# Rare microbes from diverse Earth biomes dominate community activity

**DOI:** 10.1101/636373

**Authors:** Rohan Sachdeva, Barbara J. Campbell, John F. Heidelberg

**Affiliations:** Department of Biological Sciences, University of Southern California, Los Angeles, CA 90089, USA; Innovative Genomics Institute, University of California, Berkeley, 94720, California, USA; Department of Biological Sciences, Life Science Facility, Clemson University, Clemson, SC 29634, USA; Center for Dark Energy Biosphere Investigations, University of Southern California, Los Angeles, CA 90089, USA; Wrigley Institute for Environmental Studies, University of Southern California, Los Angeles, CA 90089, USA

## Abstract

Microbes are the Earth’s most numerous organisms and are instrumental in driving major global biological and chemical processes. Microbial activity is a crucial component of all ecosystems, as microbes have the potential to control any major biochemical process. In recent years, considerable strides have been made in describing the community structure, *i.e*. diversity and abundance, of microbes from the Earth’s major biomes. In virtually all environments studied, a few highly abundant taxa dominate the structure of microbial communities. Still, microbial diversity is high and is concentrated in the less abundant, or rare, fractions of the community, *i.e*. the “long tail” of the abundance distribution. The relationship between microbial community structure and activity, specifically the role of rare microbes, and its connection to ecosystem function, is not fully understood. We analyzed 12.3 million metagenomic and metatranscriptomic sequence assemblies and their genes from environmental, human, and engineered microbiomes, and show that microbial activity is dominated by rare microbes (96% of total activity) across all measured biomes. Further, rare microbial activity was comprised of traits that are fundamental to ecosystem and organismal health, *e.g*. biogeochemical cycling and infectious disease. The activity of rare microbes was also tightly coupled to temperature, revealing a link between basic biological processes, *e.g*. reaction rates, and community activity. Our study provides a broadly applicable and predictable paradigm that implicates rare microbes as the main microbial drivers of ecosystem function and organismal health.

## Background

Members of the rare biosphere have been recognized as important drivers of many key ecosystem functions^1–6^. Rare microbes may control ecosystems as keystone members, *i.e*. community members that the whole ecosystem depends. For example, marine rhizobia are rare, but control the input of bioavailable nitrogen via N_2_ fixation^7^. Also, rare microbes may be disproportionately active relative to their abundance, *e.g*. the rarest detectable taxon by cell count in Lake Cadagno, Switzerland, an oligotrophic lake, was discovered to contribute to >40% and >70% of the total ammonium and carbon uptake^8^, respectively. However, these instances do not demonstrate the influence of rare microbes on total community activity. Sequencing of RNA transcripts and DNA of single marker genes have been employed to understand the influence of rare microbes on community activity. These methods have revealed that rare microbes are potentially more active than abundant ones^9–11^, but can suffer from over extrapolations of genome wide function and activity from a single gene^12^. Specifically, ratios of rRNA gene transcripts and rRNA gene quantities have been shown to be poor indicators of cell wide activity^13^, *e.g*. cyanobacteria can have elevated levels of rRNA in dormant cells relative to vegetative cells^14^.

To better understand the influence of rare microbes on community activity we employed a systems based approach to examine the molecular activity of the genomic content of microbiomes. We *de novo* assembled environmental DNA and RNA shotgun sequences from the genomic and transcriptomic reservoir of the global microbiome. Broadly, data are sourced from publicly available and novel environmental, host-associated, and human-engineered shotgun sequenced communities. Samples encompass the ocean, Amazon River^15^ and its plume into the ocean^16,17^, the human gut^18^, permafrost soil layers^19^, a thermokarst bog^19^, and human-engineered biogas plants^20,21^. Ocean samples span: the sunlit epipelagic^22–28^ (0 - 200 m), the dimly lit mesopelagic (200 - 1,000 m), dark bathypelagic (1,000 - 4,000 m), the benthic zone (near seafloor), and hydrothermal vent plumes^29,30^ (Table S1). Our analysis uses database independent *de novo* high resolution 99% average nucleotide (ANI) contiguous assemblies that captures the entire genomic repertoire from all domains of life. Using these assemblies, we examined the activities of rare and abundant microbes and their functional traits across many disparate environments.

## Results/Discussion

Microbial activity is consistently and commonly dominated by rare microbes. Relating community structure as a function of community activity in rare sequence assemblies across disparate environments consistently showed that total RNA expression of rare microbes was many folds higher than the total RNA expression of abundant microbes (Figs. 1, S1 – S3). Approximately 96% of all microbial activity was contributed by rare microbes (Table S2). In >90% of samples, >90% of total community activity was in the rare fraction (Fig. S4). Rare microbes were defined as those with sequence assemblies that are in the “tail” of the DNA rank abundance curve^31^, in this study >1000th rank or ~0.005% by relative abundance (Fig. S5A). Microbial activity was also dominated by rare microbes using assembly independent kmer counting (Fig. S3A), indicating that our finding is not a result of sequence assembly bias. Further, rare microbial expression was also highly overrepresented at the gene level (Fig. S3B), *i.e*. open reading frames (ORFs), and an approximate functional level based on clustering of ORFs at 60% amino acid identity (Fig. S3C). Sense, or coding strand, RNA transcript expression of ORFs was higher than antisense expression for rare microbes (Mann-Whitney *U* test; *P* < 2.2 10^−16^) that were from samples prepared with methods that retained strand orientation (Figs. 2D and S6). Sense mRNAs are transcripts that are meant for downstream translation into protein, whereas antisense transcripts primarily act as post-transcriptional regulators by directly binding to sense transcripts^32^. Higher sense transcription indicates that most microbial transcription is ultimately meant for protein translation. This pattern is consistent across all sampled environments and lifestyles, *e.g*. free-living or attached (Fig. 2E). Further, our analysis captures the entire genomic representation of microbial communities and is not limited to single genes or processes.

**Figure 1.**
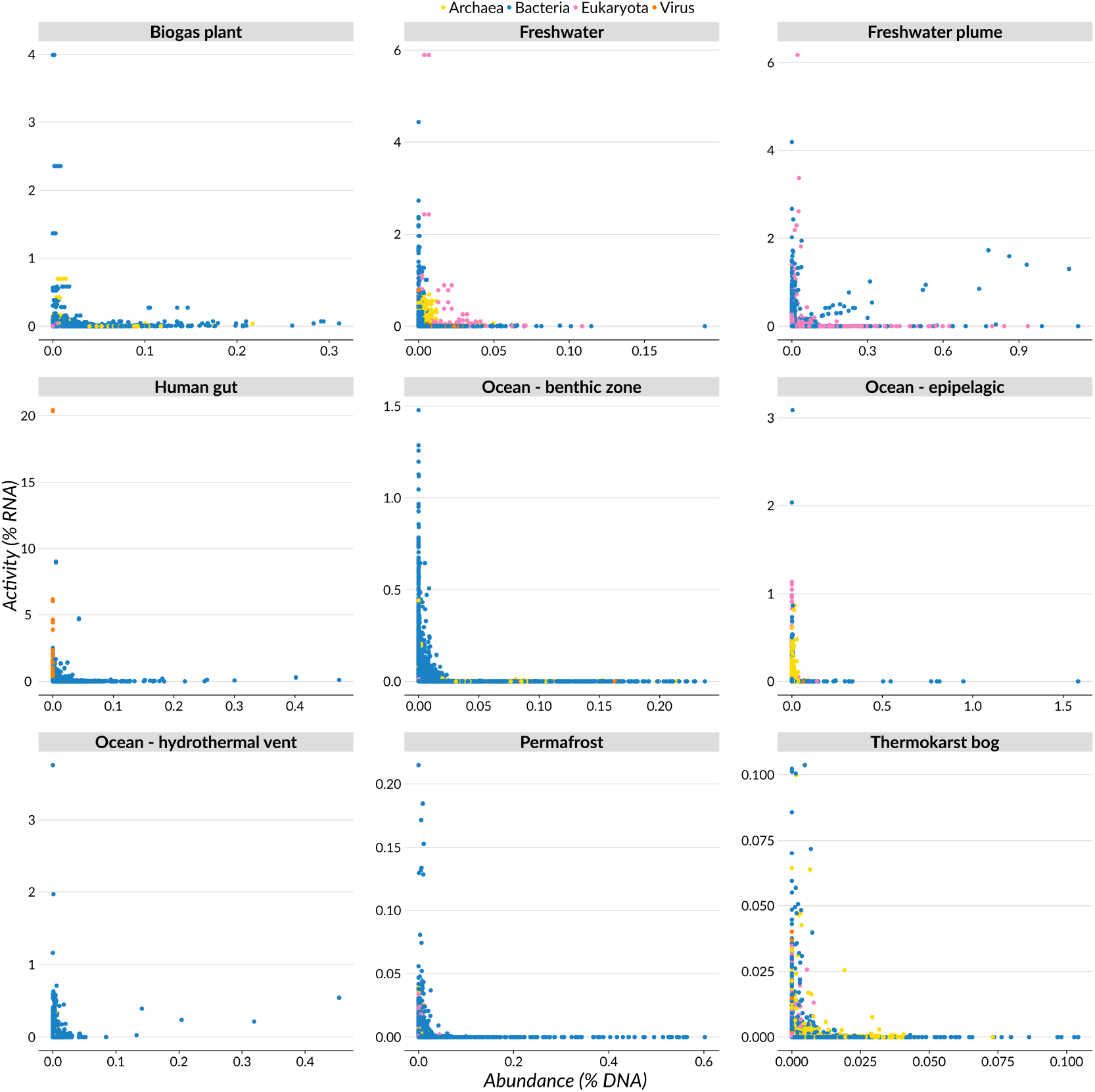
Relationships between community structure and community activity of highly resolved sequence assemblies across disparate environments. Each point represents a sequence assembly and its relative contribution to community structure and activity. Environments are biogas plants, freshwater, freshwater plume into the ocean, the human gut, ocean (epipelagic, benthic zone, hydrothermal vents), permafrost, and thermokarst bog. Community structure is expressed as relative frequencies of DNA and community activity as relative frequencies of RNA of samples sequenced with Illumina platforms. All frequencies were adjusted to sequence assembly length and subsampled to account for uneven sequencing effort. Points are colored by RefSeq lowest common ancestor (LCA) taxonomy at the domain rank.

**Figure 2.**
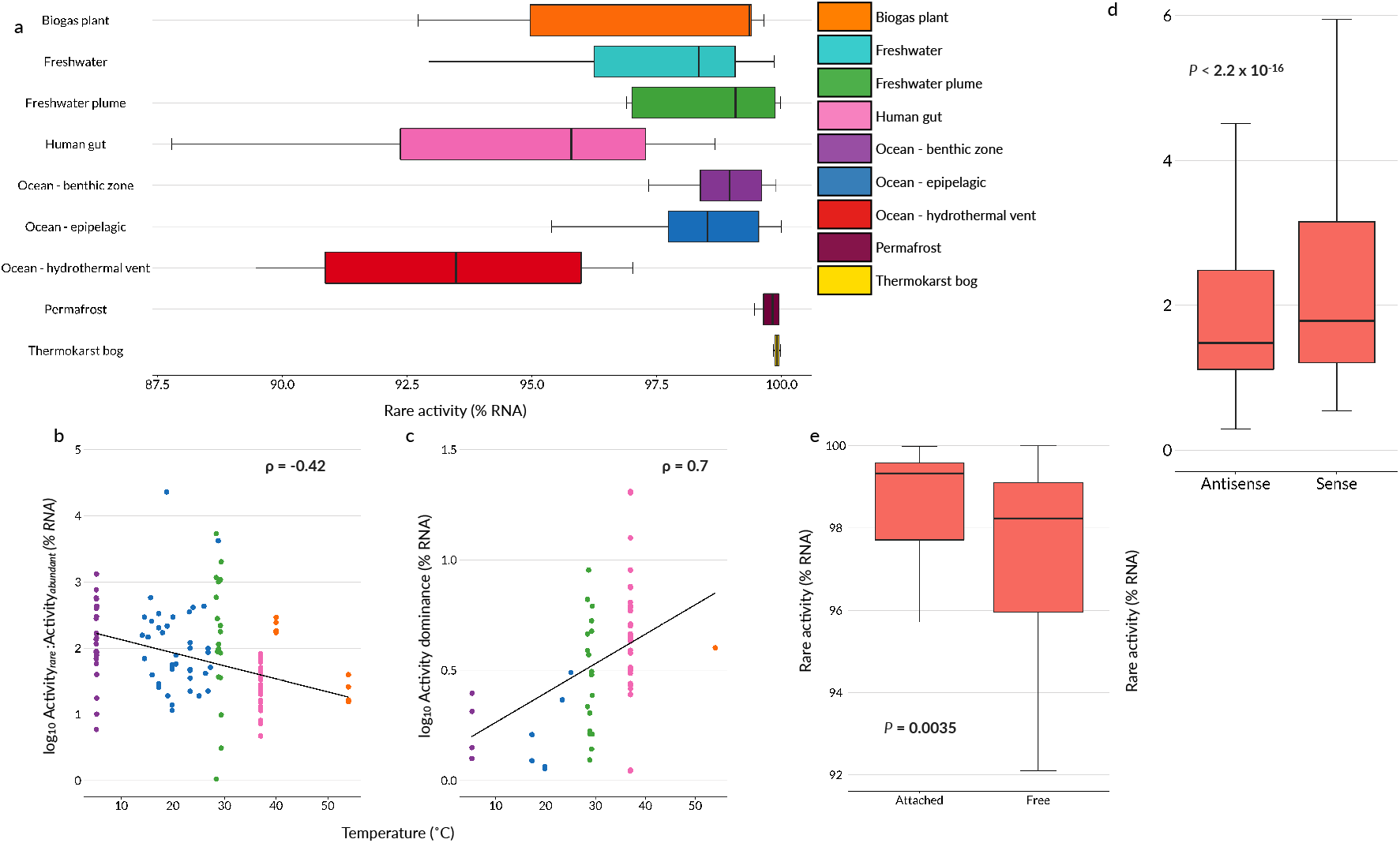
Patterns of rare high-resolution sequence assembly activity of Illumina sequenced samples across different environments, lifestyles, and temperatures. **a)** Box and whisker plots of rare assembly activity across different environments. Mann–Whitney *U* test; *P* < 2.2 x 10^−16^. **b)** Rare microbial activity expressed as a ratio of log10 rare:abundant % RNA as a function of temperature. Linear regression plotted with Spearman’s ρ = −0.42 and P = 1.701 x 10^−7^. **c)** Activity dominance expressed as Berger–Parker dominance of % RNA as a function of temperature. Linear regression plotted with Spearman’s ρ = −0.7 and *P* < 2.2 x 10^−6^. **d)** Box and whisker plots of rare antisense and sense microbial activity. **e)** Box and whisker plots of rare microbial activity in attached and free, *i.e*. planktonic, lifestyles. Samples with microbes with attached lifestyles were considered: sediment samples, *i.e*. permafrost, thermokarst bog, and aquatic (freshwater and marine) samples collected on filters >0.8 μm. Samples with microbes with free lifestyles were considered: the human gut and aquatic (freshwater and marine) samples collected on filters 0.1 - 3.0 μm.

Next, we examined the structure of microbial abundance and activity across environments. Samples clustered by environment across DNA, RNA, and specific activity^10^ (RNA:DNA) distributions (Fig. S7). Further, the rank abundance distributions, *i.e*. measures of biodiversity, from all environments using sequence assemblies follow a highly skewed curve, as has been widely reported for microbial communities using single-marker genes^5,31,33^ (Fig. S5A). This demonstrates that highly skewed rank abundance curves even at a genomic level are a consistent feature of microbial communities regardless of environment. Highly skewed rank abundance curves suggest that there are dominant genotypes in microbial populations, despite potentially high recombination rates^34^. The prevalence of dominant genotypes indicate that ecological pressures, *e.g*. nutrient limitation and predation, can select for specific “winning” genotypes among highly related microbial populations. RNA activity patterns were similarly skewed by rank expression, demonstrating that microbial activity is similarly dominated by a small number of overrepresented assembled sequences (Fig. S5B). Community activity was also more skewed toward a few assemblies, as indicated by lower Pielou’s evenness (*J*’) compared to community structure *J*’ (Fig. S8; Mann–Whitney *U* test; *P* < 2.2 x 10^−16^). The rank abundance and rank activity curves reflect that not only do a small number of microbes dominate community structure, but fewer types, relative to community structure, also dominate community activity.

The extent that rare microbes contributed to activity varied within and between environments (Fig. 2A). This pattern was driven by temperature variation, as total activity of rare microbes decreased with increasing temperature (Fig. 2B; Spearman’s ρ = −0.42; *P* = 3.979 x 10^−7^). The degree that community activity was dominated by fewer microbes, *i.e*. activity dominance, alternatively increased with increasing temperature (Fig. 2C; Spearman’s ρ = 0.7; *P* < 2.2 x 10^−16^). Temperature is a first order determinant of chemical reaction kinetics, and, therefore, biochemical processes^35^. Higher temperatures induce higher metabolic rates that ultimately mediate biological activities^36^. This positive relationship between temperature and biological process rates has been implicated in controlling many ecological^37–39^ and evolutionary^40,41^ patterns. Our observations show that temperature is also a major mediator of the structure of microbial community activity.

We explored the functional contribution of rare microbes to community activity across all environments. Functional traits were examined that were more than two fold overexpressed in the rare fraction relative to the abundant fraction (Mann–Whitney *U* test; *P* < 3.9 x 10^−71^). Rare ORFs were enriched in the activity of functional traits that have a direct influence on total ecosystem function (Fig. 3), *e.g*. energy production and conversion, carbohydrate transport and metabolism, coenzyme transport and metabolism, inorganic ion transport and metabolism, nucleotide transport and metabolism, amino acid transport and metabolism, lipid transport and metabolism. Rare ORFs were also enriched for functional traits involved with cell motility, coinciding with the observation that rare microbes tend toward chemotactic lifestyles^42–44^ (Fig. 3A). Other processes linked to growth were also overrepresented in rare ORFs, including cell growth and death, cell cycle control, cell division, chromosome partitioning, cell wall/membrane/envelope biogenesis, and translation (Fig. 3AB). This suggests that rare microbes have higher growth rates, as well as the aforementioned higher metabolic activity. Although higher growth rates should result in higher abundance, rare microbes were enriched in defense mechanisms and xenobiotic degradation, suggesting that they are subject to higher pressures of viral predation, grazing, host-defense, and allelopathy (Fig. 3B). Rare ORFs were also enriched in infectious disease categories, implicating rare microbes in animal and human disease. Finally, many of the genes most expressed by rare microbes were directly involved in major biogeochemical processes, such as photosynthesis, N_2_ fixation, and ammonia oxidation (Fig. 3C). For example, a rare *Candidatus* Atelocyanobacterium thalassa^45^ (unicellular cyanobacteria group A member) had a *nifH* (iron binding component of nitrogenase) with the highest annotatable contribution to activity in the epipelagic South Atlantic Ocean (Table S3). Other relevant biochemical processes include ammonium, phosphate, and energy driven carbohydrate ABC transport. The influence of rare microbes in mediating important ecosystem processes highlights their role as keystone members of ecosystems.

**Figure 3.**
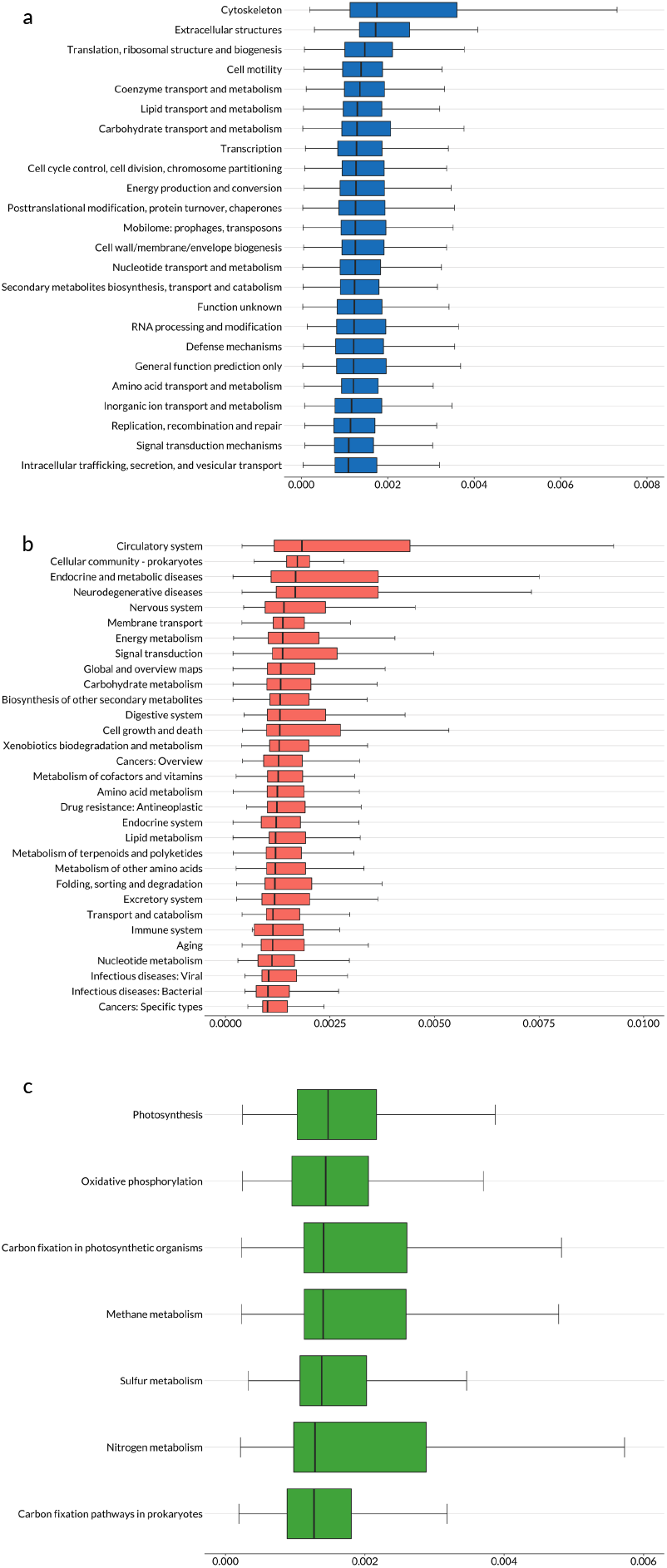
Functional traits of Illumina sequenced rare microbial activity enriched in rare fraction relative to the abundant fraction. Overexpression of traits in the rare fraction was determined by selecting ORFs with >2X activity in the rare fraction that had a Mann–Whitney U with *P* < 1 x 10^−70^ **a)** Box and whisker plots of rare overexpressed NCBI Clusters of Orthologous Groups (COGs) functional categories. **c)** Box and whisker plots of KEGG BRITE categories overexpressed in the rare fraction. **d)** Box and whisker plots of KEGG BRITE “Energy metabolism” subcategories overexpressed in the rare fraction. Box and whisker plots are sorted by median and exclude ORFs where % RNA was 0.

Deciphering the role of rare microbes in microbial communities is at the core of understanding the influence that microbes have on ecosystem function. We demonstrate that rare microbes are not only more active, but dominate microbial community activity in many different environments. This pattern is consistent across many disparate environments, ranging from the human gut to human engineered biogas plants to hydrothermal vents. The contribution of rare microbes to community activity varies across environments and is strongly influenced by temperature, implying that fundamental biological processes, *e.g*. reaction rates, control community activity structure. Rare microbes were more involved than abundant microbes in important ecosystem processes, *e.g*. energy transformation and biogeochemical cycling. Rare microbial activity was also enriched in genes related to infectious disease, underscoring the detrimental impacts of rare microbes on animal and human health. Our observations indicate that the dominance of rare microbial activity is a conserved trait for all of Earth’s biomes.

## Methods

### Sargasso Sea sample collection and sequencing

Four samples were collected from the end of spring (3/24/2010 and 3/26/2010) and summer (8/20/2010 and 8/22/2010) from the Bermuda Atlantic Time Series station (BATS, 31° 29’ 46” N, 63° 59’ 52” W). 5 - 20 L were collected from a depth of 50 m and immediately amended with an equal volume of RNAlater once on board ship. The RNAlater amended samples were sequentially filtered through a glass fiber 0.8 μm GF/F filter (Whatman) and finally onto a 0.22 μm Durapore (Millipore) filter within one hour of collection and stored at −80 °C. DNA and RNA were extracted as previously described in Campbell *et al*. 2009^46^. DNA was fragmented with a Covaris S2 with the recommended parameters to generate 300 bp inserts prior to metagenome library preparation using an Encore NGS Library System I (NuGEN). RNA was purified from DNA as described previously^46^ and approximately 1 μg from spring samples were rRNA subtracted using the MICROBExpress Bacterial mRNA Enrichment Kit (Ambion). All RNAs were reverse transcribed following the manufacturer’s instructions for cDNA synthesis and metatranscriptome libraries prepared with an Ovation RNA-Seq System (NuGEN) kit. Libraries were sequenced via a paired end 2 x 100 bp strategy using an Illumina HiSeq 2000. Libraries from DNA extracted from 3/24/2010 and 8/20/2010 samples were also sequenced on the Roche 454 GS FLX+ platform and using circular consensus sequencing on a PacBio RS.

### San Pedro Ocean Time-series (SPOT) benthic zone sample collection and extraction

Seawater samples (1 - 20 L) were collected approximately monthly over ~2.5 years (n = 32) from near the ocean floor (890 m) in the Southern California Bight at the SPOT station (33° 33’N, 118° 24’W) aboard the *R/V Yellowfin*. Samples were serially filtered through a nylon mesh (80 μm), Acrodisc (Pall) glass fiber filter (~1 μm), and terminally with a Sterivex-GP (Millipore) polyethersulfone filter (0.22 μm). Acrodisc and Sterivex filters were preserved with 250 μl and 500 μl of RNAlater, respectively, and subsequently incubated at ambient temperature for 2 min. Samples were immediately frozen in LN_2_ and stored at −80°C. The 0.22 μm - 1μm fraction was selected for sequencing and DNA was extracted from each Sterivex cartridge using a modified AllPrep DNA/RNA (Qiagen) kit. RNAlater was sparged from each Sterivex using a syringe. High salt concentrations can lower DNA yields with the AllPrep DNA/RNA kit. To desalt RNAlater while maximizing nucleic acid concentrations we used an Amicon Ultra 3k. Briefly, 500 μl RNAlater (Ambion) was transferred to an Amicon Ultra 3k 0.5 ml spin concentrator and centrifuged at 14,000 g for 30 min at 4°C. This was repeated with any remaining unconcentrated RNAlater. 400 μl of 4°C nuclease free water was added directly to the concentrator and concentrate to desalt and centrifuged at 14,000 g for 1 hr at 4 °C 2 ml of 65°C RLT+ lysis buffer and 20 μl beta-mercaptoethanol were added to the previously desalted and concentrated RNAlater. The Sterivex cartridge was sealed with luer lock caps and vortexed for 5 s, flipped and vortexed again for 5s. The Sterivex cartridge was horizontally rotated for 15 min at 65°C. The lysate was removed, and the Sterivex filter was lysed again with 1ml RLT+ and 10 μl beta-mercaptoethanol. The lysates were combined and DNA was purified following the manufacturer’s protocol and finally eluted with buffer AE. RNA was purified from the DNA purification flow through following the manufacturer’s instructions with a 30 min on-column DNAse step and a final elution with 50 μl nuclease free water. 1 μl of RiboGuard RNase Inhibitor (Epicentre) was added to the eluted RNA to protect against ribonucleases. RNA quantities were determined using a Qubit RNA High Sensitivity (ThermoFisher) kit.

### SPOT benthic zone metagenome library construction and sequencing

DNA concentrations were determined for each sample using a Qubit High Sensitivity DNA assay (Invitrogen) and diluted to 10 ng. Samples were amended with the DNA of 4 exotic genomes from American Type Culture Collection with a range of 34.5 - 59.9 %G+C content (Table S4). Each genome was added in 2 fold increasing concentrations at ~1% of the total DNA concentration of each sample. Metagenomes were prepared from each amended sample using a Covaris S2 (130 μl) with parameters: duty cycle (10%), intensity (5) cycles/burst (200), time (60 s). Insert and dual indexed libraries were prepared using a modified NEBNext Ultra DNA II dual indexing kit (New England Biolabs) without size selection. The protocol was modified to control for chimeric PCR amplicons and reduce overamplification biases. Adaptor ligated and end repaired fragments were amplified with 0.1X of 10,000X SYBR Green I (Invitrogen). Amplification was monitored on a CFX96 real-time PCR machine (BioRad) and stopped in the exponential phase to avoid overamplification. The reaction was held at 98°C for 30 s followed by 6 cycles of amplification at 98°C for 10 s, 65°C for 5 min, and a final extension of 30 min. Extension times were increased to reduce chimeric amplicons. Libraries were subsequently 2 x 250 bp paired end sequenced on an Illumina HiSeq 2500.

### SPOT benthic zone metatranscriptome library construction and sequencing

40 ng of RNA was amended with 8 μl of a 10,000X dilution of External RNA Control Consortium^47^ (ERCC) Spike-In Mix 1 (Ambion). The ERCC mix consists of 92 transcripts ranging from 250 to 2,000 nt in length with a large dynamic fold range. The transcripts are mostly novel synthesized sequences, but do contain *Bacillus subtilis* transcripts. The added genome and ERCC controls are sequencing controls used to verify that our sample preparation and bioinformatic protocols are accurate and reproducible. The amended RNA samples were rRNA depleted using a RiboZero Bacteria rRNA removal kit. RNA was fragmented using a Covaris S2 targeting 600 bp: sample volume (130 μl), peak incident power (50 W), duty factor (20%), cycles per burst (200), treatment time (60 s), temperature (7°C). Strand specific cDNA was generated using random priming without size selection from a NEBNext Ultra RNA directional kit. Illumina libraries were constructed from the resulting cDNA using a modified NEBNext DNA Ultra II dual indexing kit. The protocol was modified to use RT-PCR to control for overamplification and longer extension times to control for chimeric amplicons as previously described (Chapter 1), and resulted in 12 cycles of amplification. Metagenomic and metatranscriptomic libraries were sequenced 2 x 250 bp using an Illumina HiSeq 2500.

### Sequence and metadata retrieval

Raw metagenomic and metatranscriptomic reads from Illumina, 454, Sanger, and PacBio sequencing platforms from 504 sequence libraries covering a number of disparate environments were downloaded for each publicly available dataset (Table S1). Ocean samples encompass the epipelagic, mesopelagic, bathypelagic, the benthic zone, and hydrothermal vent plumes, ranging in depths 0 - 4,946 m. Epipelagic samples include coastal and open ocean sites. Other samples come from the Amazon River and its plume into the Atlantic Ocean, active and frozen permafrost, a thermokarst bog, human guts, and biogas plants. Data were retrieved from iMicrobe, NCBI SRA, and ENA (Table S1). Metagenome and metatranscriptome sample pairings and sample metadata (*e.g*. temperature and environment) were determined using sequence database metadata or directly from the source publications. Human gut sample temperatures were not publicly available and were inferred to be 37°C based on the typical human body temperature^48^.

### Sequence quality control and assembly

The majority of samples were sequenced using Illumina based flow cell technologies (Table S1). Illumina reads were adapter and quality trimmed in one pass to retain the largest regions with Q > 25 (BBMap^49^ v36.19; bbduk.sh qtrim=rl trimq=25 ktrim=r k=25 mink=11 hdist=1 ref=truseq.fa.gz). Reads from long read sequencing platforms, Roche 454, Sanger, and Pacbio RS, were similarly trimmed to Q > 20 (BBMap v36.19; bbduk.sh qtrim=rl trimq=20). Genome and ERCC controls were removed from each SPOT benthic zone sample prior to assembly and mapping. Reads were mapped to the added genome and ERCC controls at 95% identity and the unmapped reads and their paired end mates were retained for downstream assembly and mapping (BBMap^49^ v36.19; bbmap.sh idfilter=0.95). Each metagenome and metatranscriptome from each sample was individually assembled *de novo* using MEGAHIT^50^ in paired end mode if sequence libraries were paired end sequenced. MEGAHIT was run to ensure that no bubbles were merged that were < 99% to ensure that only highly-resolved assemblies were generated (MEGAHIT v1.0.6; megahit --merge-level 20, 0.99 --k-min 21 --k-max 255 --k-step 6). All assemblies were combined and dereplicated using a semi-global alignment method that merged together assemblies that were totally contained and >99% similar (BBMap v36.19; dedupe.sh minidentity=0.99). Assemblies <1 kb were discarded. Processing resulted in 12,338,658 assemblies, comprised of 25.48 Gbp.

### Annotation

Open reading frames (ORFs) were predicted (ORFfinder^51^ -s 1 -n T) and ORFs > 200 amino acids were retained. The resulting ORFs were searched against the KEGG reference database (DIAMOND^52^; diamond blastp -e 1e-5 --sensitive), KEGG modules and pathways were retrieved using the KEGG API. Only the best hits with E-value < 1 x 10^−10^ were assigned. ORFs were also searched against the complete non-redundant NCBI RefSeq Release 83^53^ protein database (DIAMOND; diamond blastp --top 5 -e 1e-5) and hits with E-value < 1 x 10^−10^ were retained. A best hit was determined by sorting by E-value, bit score, and percent identity. Taxonomy was assigned for each sequence assembly using a lowest common ancestor (LCA) approach. RefSeq hits for each ORF that were within 5% of the bit score of the best hit were retained. The remaining hits were used to assign a LCA taxonomy for each assembly.

### Metagenome and metatranscriptome mapping and counting

Mapping and counting were performed within each sequencing platform where both metagenomes and metatranscriptomes were sequenced, *i.e*. Roche 454 and Illumina platforms. Small subunit (16S and 18S) and large subunit (5S, 5.8S, 23S, and 28S) rRNAs were removed from each read set prior to mapping reads from each environment to each dereplicated set of assemblies (SortMeRNA^54^ v2.1; sortmerna --paired_out --fastx --ref silva-bac-16s-id90.fasta silva-arc-16s-id95.fasta silva-euk-18s-id95.fasta rfam-5s-database-id98.fasta rfam-5.8s-database-id98.fasta silva-arc-23s-id98.fasta silva-bac-23s-id98.fasta silva-euk-28s-id98.fasta). The resulting rRNA filtered reads from each library were mapped to the dereplicated set of assemblies. Filtered reads were mapped to retain all sites with the highest score, ie. the assemblies that are the best matches, and >99% identity (BBMap v36.19; bbmap.sh ambiguous=all maxsites=1000000000 maxsites2=1000000000 sssr=1.0 secondary=t minid=0.98 idfilter=0.99).

Mapped counts were determined for each sample and all mapped sites with a minimum read length of 50 bp were considered (featureCounts^55^ v1.5.3; featureCounts -f -O -M --minOverlap 50 -s 0). ORF expression and abundance were similarly counted by using the previously mapped reads and ORF start and stop locations for counting (featureCounts v1.5.3; featureCounts -f -O - M --minOverlap 50 -s 0). Strand specific, *i.e*. sense and antisense, counts were determined for RNA libraries prepared to retain strand information (Table S1) using featureCounts v1.5.3 in forward (featureCounts v1.5.3; featureCounts -f -O -M --minOverlap 50 -s 1) and reverse (featureCounts v1.5.3; featureCounts -f -O -M --minOverlap 50 -s 2) strand counting modes. All strand specific libraries were prepared using methods that result in sequencing reads that are the reverse complement of the transcribed RNA sequence. Accordingly, the reverse counts were considered as the sense counts and the forward counts as the antisense counts. Next, assemblies that matched to the Centrifuge^56^ NCBI nucleotide index^56^ (12/06/2016) and matches to human sequences >90% assemblies coverage were removed (Centrifuge v1.0.3-beta^56^; default parameters). Assemblies that matched to the genome, ERCC, and PhiX 174 controls were also removed (NCBI BLAST+^57^ v2.6.0; blastn -evalue 0).

Prior to counting, assemblies matching mapped counts were divided by assembly length or ORF length to account for higher recruitment of longer assemblies or ORFs. Assemblies and unstranded ORF length adjusted counts were subsampled, *i.e*. rarefied, to the length adjusted counts of the smallest sample to account for uneven sequencing coverage. Stranded forward and reverse length adjusted counts were subsampled to the total forward and reverse length adjusted counts of the smallest sample. Subsampling was done separately for Illumina and 454 sequences to maintain high counts for Illumina samples because the 454 based sequencing effort was much lower than the Illumina based sequencing effort. Length adjusted and subsampled counts for each assembly or ORF were normalized to the total length normalized count of each sample to obtain a relative count, *e.g*. relative DNA abundance or relative RNA expression. Pairings between RNA and DNA counts were determined from sample database metadata or from the sample publications. In some cases, there were multiple pairings with a sample. Some samples had > 1 DNA or RNA library sequenced. For example, if a sample has 2 DNA and 2 RNA sequence libraries, there are 4 possible DNA and RNA pairs. All data manipulations were done using Python^58^ and Pandas^59^. Specific activity was determined as a ratio by dividing relative RNA and relative DNA counts^10^.

### SPOT benthic zone metagenomic and metatranscriptomic controls

Genome (DNA) and ERCC (RNA) controls were analyzed using the same mapping and counting protocols that we used to quantify abundance and expression. Mapping and counting were performed exactly as for mapping and counting of all Illumina sequences from all environments. Trimmed and rRNA filtered metagenomic and metatranscriptomic from the SPOT benthic zone samples were mapped to the genome and ERCC control sequences (BBMap v36.19; bbmap.sh ambiguous=all maxsites=1000000000 maxsites2=1000000000 sssr=1.0 secondary=t minid=0.98 idfilter=0.99). Counts (featureCounts v1.5.3; featureCounts -f -O -M --minOverlap 50 -s 0) were transformed into relative counts by length dividing and normalizing to the total mappings from each sample. Performance was quantified by plotting against the expected genome and ERCC control concentrations. Both genome and ERCC controls had high agreement between input quantities and measurements (R^2^ > 0.99) using the afforementioned protocols (Fig. S9).

### Approximate functional clusters

Functional clusters were generated by clustering ORFs based on percent identity. ORFs longer than 200 amino acids were grouped into protein clusters with >60% identity (cd-hit^60^ v4.6; cd-hit -c 0.6 -n 4). Relative counts for clusters were determined by summing the length adjusted subsampled normalized counts of each ORF contained within a cluster.

### Assembly free kmer based analysis

Two samples were randomly selected from each environment for verification of abundance and activity relationships using an assembly free kmer counting. Counts of all canonical 31 letter kmers were generated for each sample (BBMap v36.19; kmercountexact.sh k=31). Kmers of 31 letters were chosen because they have been shown to be able to distinguish approximate microbial species^61^. Kmers with low complexity (Shannon entropy ≤ 1.84) were removed to increase the likelihood of capturing kmers that can separate microbes at the species-level.

### Diversity metrics and statistics

Diversity metrics were calculated using relative counts for metagenomes and metatranscriptomes. Pielou’s evenness (*J*’) and Berger–Parker dominance were calculated using scikit-bio. Linear regressions and nonparametric statistics were calculated using Spearman’s ρ and the Mann-Whitney U test from the R^62^ stats package. Non-metric dimensional scaling plots (NMDS) were generated using the vegan^63^ R package and Bray-Curtis dissimilarities (vegan v2.4-4; metamds (distance=“bray”).

## Acknowledgements

We would like to acknowledge the Wrigley Institute for Environmental Studies, the crew of the *R/V Yellowfin*, and Troy Gunderson for logistical support of SPOT samples. We thank Jed Fuhrman and his lab, including Erin Fichot, Catherine Garcia, David Needham, Alma Parada, Ella Sieradzki, Jacob Cram, and Vicki Trinh. We also thank Megan Hall for helpful comments on drafts of the manuscript. We thank the scientists who deposited their sequences in publicly accessible databases. This work was funded by the Wrigley Graduate Summer Fellowship Program and National Science Foundations awards 12053, 0824981, and 0939564 to J.F.H., and 0825468 and 1261359 to B.J.C.

## Contributions

R.S., B.J.C, and J.F.H designed the study for epipelagic and benthic marine environments. R.S. expanded the study to other environments. R.S. prepared SPOT benthic zone metagenomes and metatranscriptomes. B.J.C prepared Sargasso Sea metagenomes and metatranscriptomes. R.S. analyzed the data. R.S wrote the paper with input from B.J.C and J.F.H. All authors read and approved the manuscript.

## Data availability

Sargasso Sea sequences are deposited under NCBI SRA BioProject PRJNA242360. SPOT sequences are currently being deposited at NCBI.

## Supplementary information

**Figure S1.**
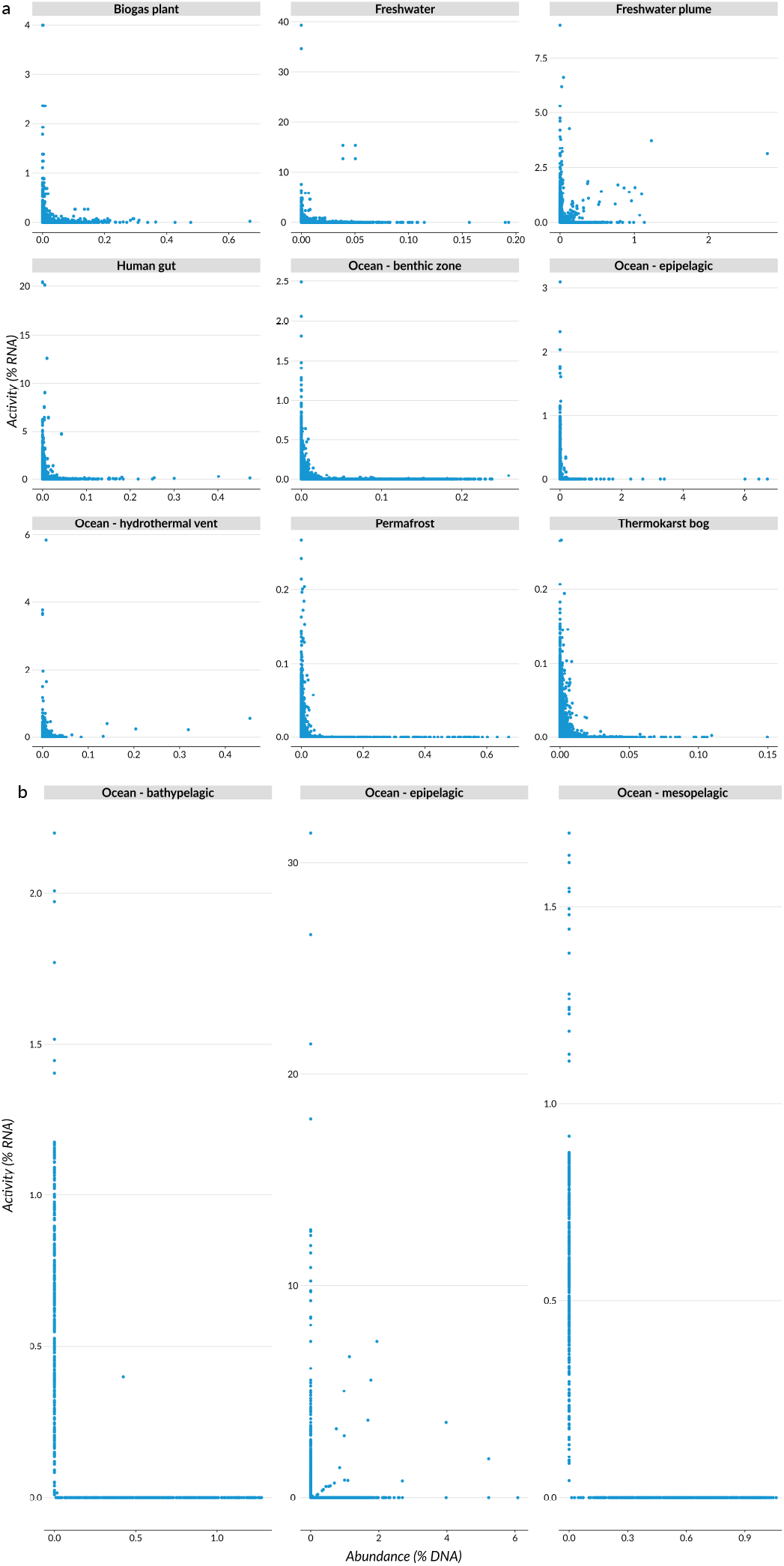
Relationships between relative abundance (% DNA) and activity (% RNA). Relationships between relative abundance and activity regardless of taxonomy that were sequenced with the a) Illumina and b) Roche 454 platforms.

**Figure S2.**
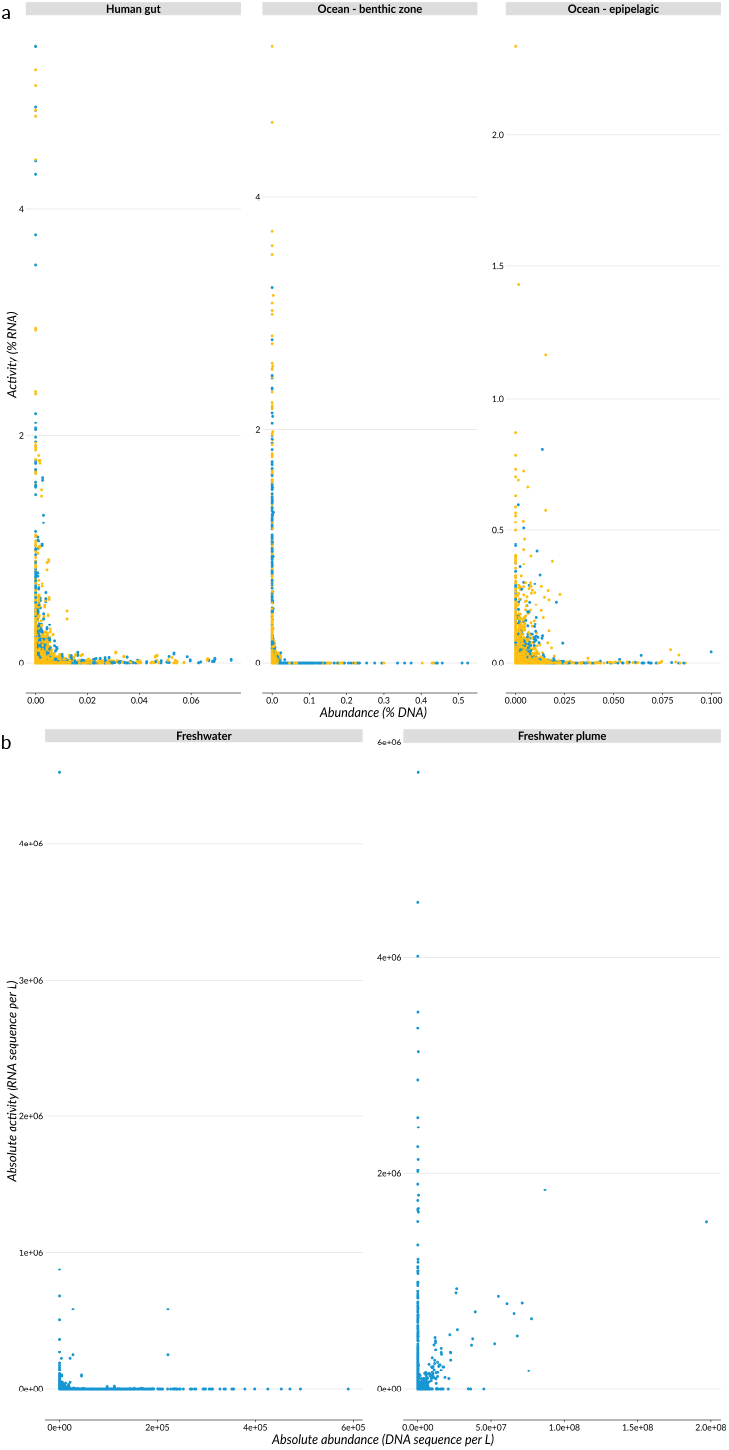
Relationships between abundance and activity for samples sequenced with methods that retain RNA strand direction and absolute quantifications. a) Stranded (directional) relationships between relative abundance and relative activity. Strandedness is only retained at the ORF level therefore mappings were counted at the ORF level. The sense (yellow) reflects transcripts that directly correspond to mRNA that will be translated into peptides. Antisense (blue) transcripts directly correspond to transcripts that are the reverse complement of mRNA and can act as transcriptional and post-translational regulators^1^. b) Relationships between absolute abundance (DNA sequence L^−1^) and absolute activity (RNA sequence L^−1^) from the Amazon River^2^ and the Amazon River plume^3,4^. Absolute quantities were generated by multiplying relative frequencies by measured quantities of DNA and RNA from Satinsky et al., 2014a^19^, Satinsky et al., 2014b^20^, Satinsky et al., 2015^18^. Briefly, DNA and RNA samples were amended with known quantities at the time of nucleic acid extraction. The percent recovery of the DNA and RNA additions provide a direct conversion of relative frequencies.

**Figure S3.**
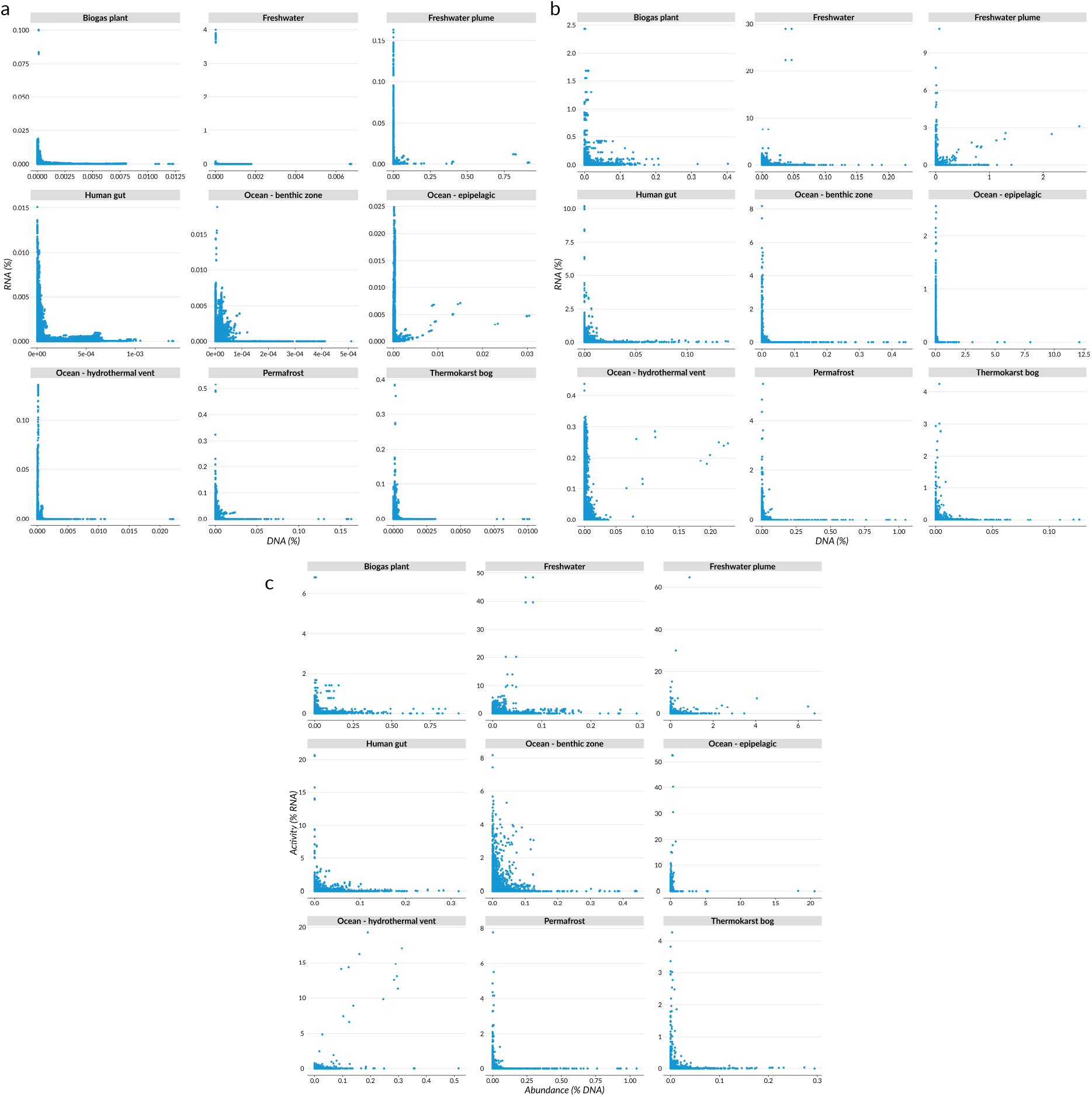
Relationships between relative abundance (% DNA) and relative activity (% RNA) of kmers, ORFs, and approximate functional clusters. a) Abundance and activity of microbial communities using kmer counts. Kmers with low complexity were removed and only those with counts >2 per sample that were represented in both DNA and RNA fractions were retained. b) Relationships between abundance and activity of ORFs. c) Abundance and activity of functional protein clusters were generated by semi-global clustering of ORFs at 60% identity. Relative frequencies for each cluster were generated by summing the relative frequencies of ORFs that formed each cluster.

**Figure S4.**
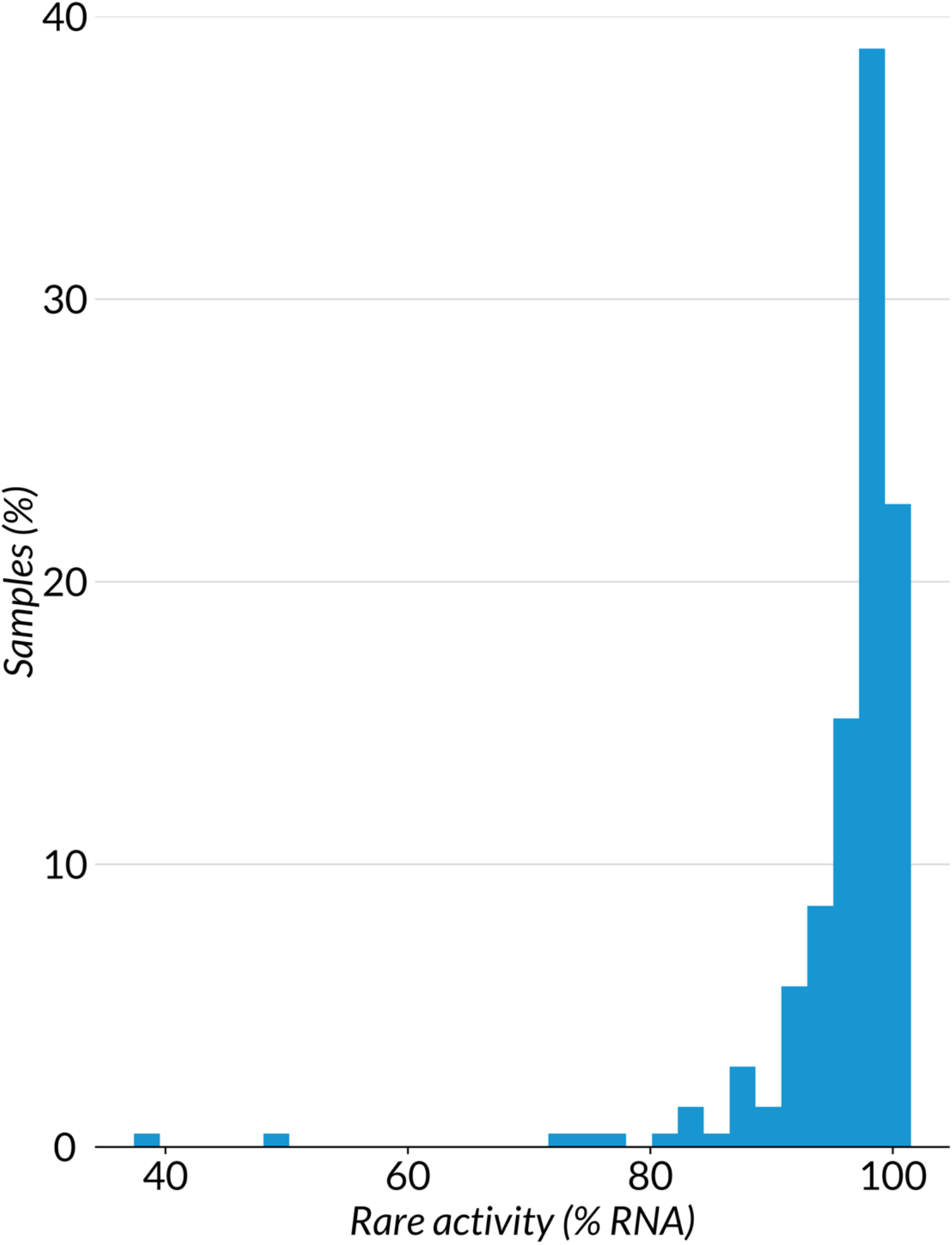
Distribution of rare activity across all Illumina sequenced samples. Rare activity is expressed as the relative RNA frequencies of assemblies in the rare fraction.

**Figure S5.**
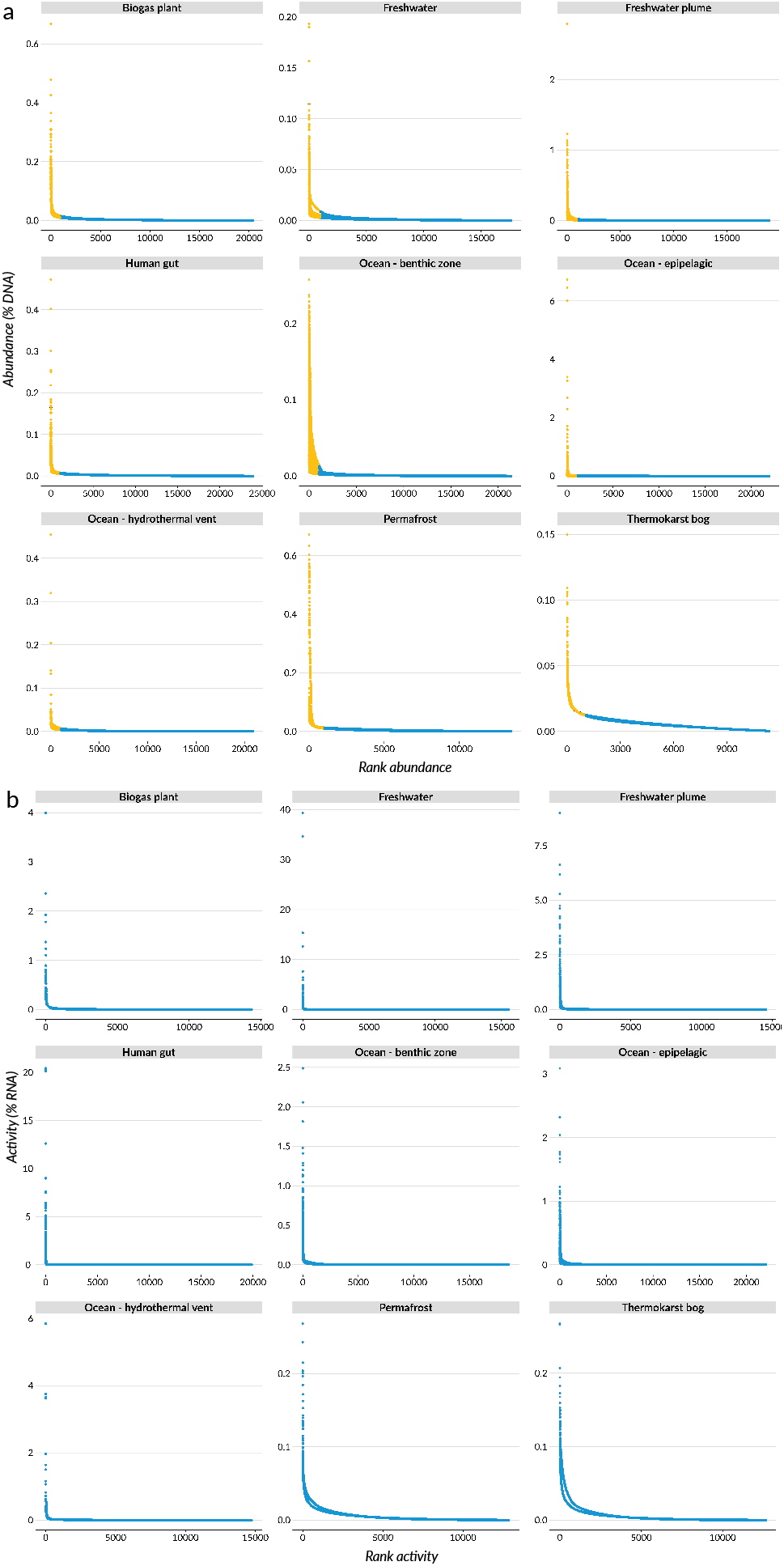
Relative abundance (% DNA) as a function of DNA rank abundance of metagenomic and metatranscriptomic sequence assemblies for Illumina sequenced samples. a) Rank abundance curves by environment. Rare assemblies are colored blue and abundant assemblies are yellow. Rare is considered >1000th rank and are in the tail of the rank abundance curves or a mean of 0.005%. b) Rank activity curves by environment. Rank activity is plotted as activity (% RNA) as a function of rank activity.

**Figure S6.**
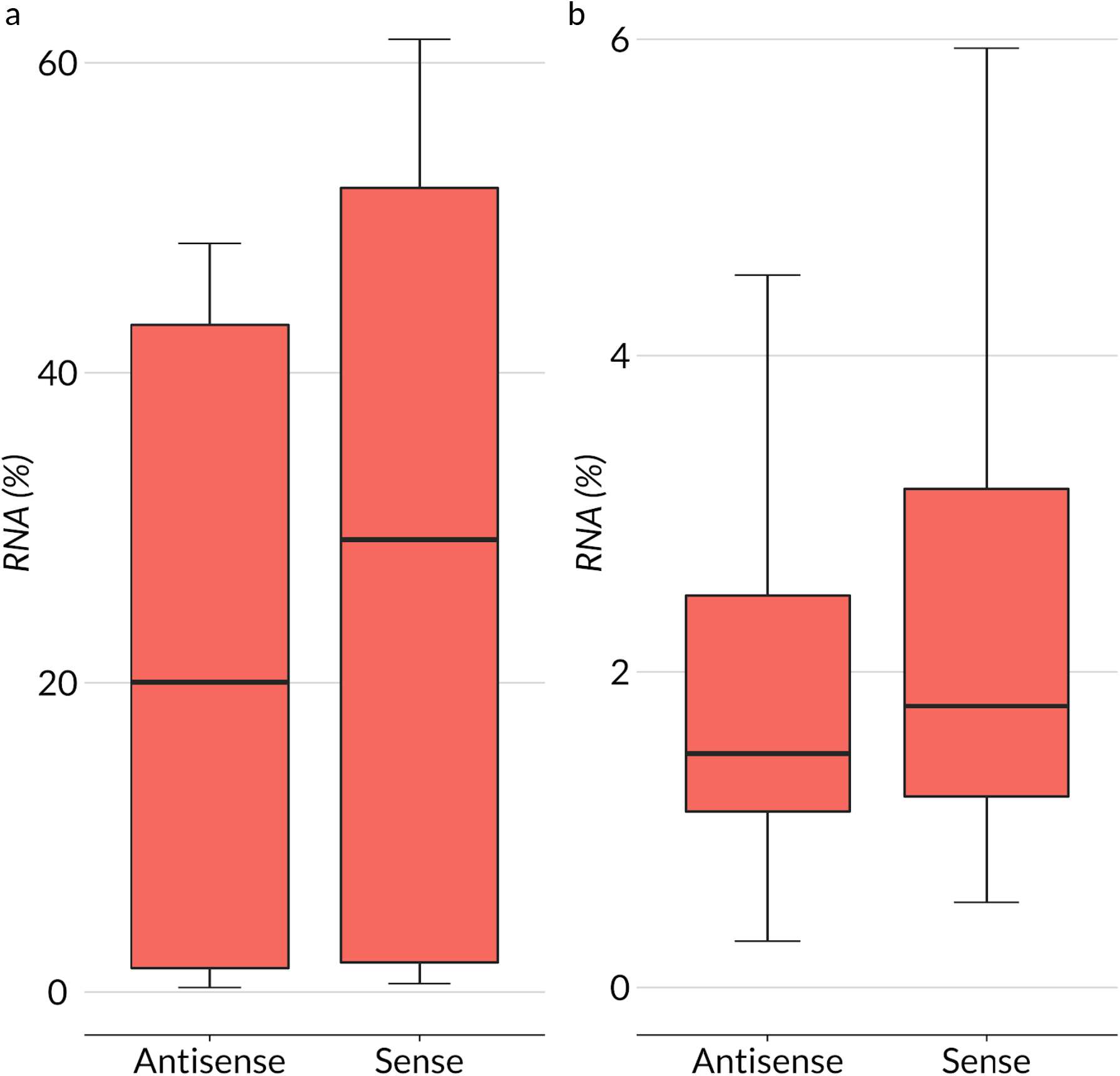
Comparison between antisense and sense expression in whole and rare community fractions. a) Box and whisker plot of antisense and sense rare microbial activity (*P* = 3.6 x 10^−6^). b) Box and whisker plot of antisense and sense whole community activity (*P* = 0.1).

**Figure S7.**
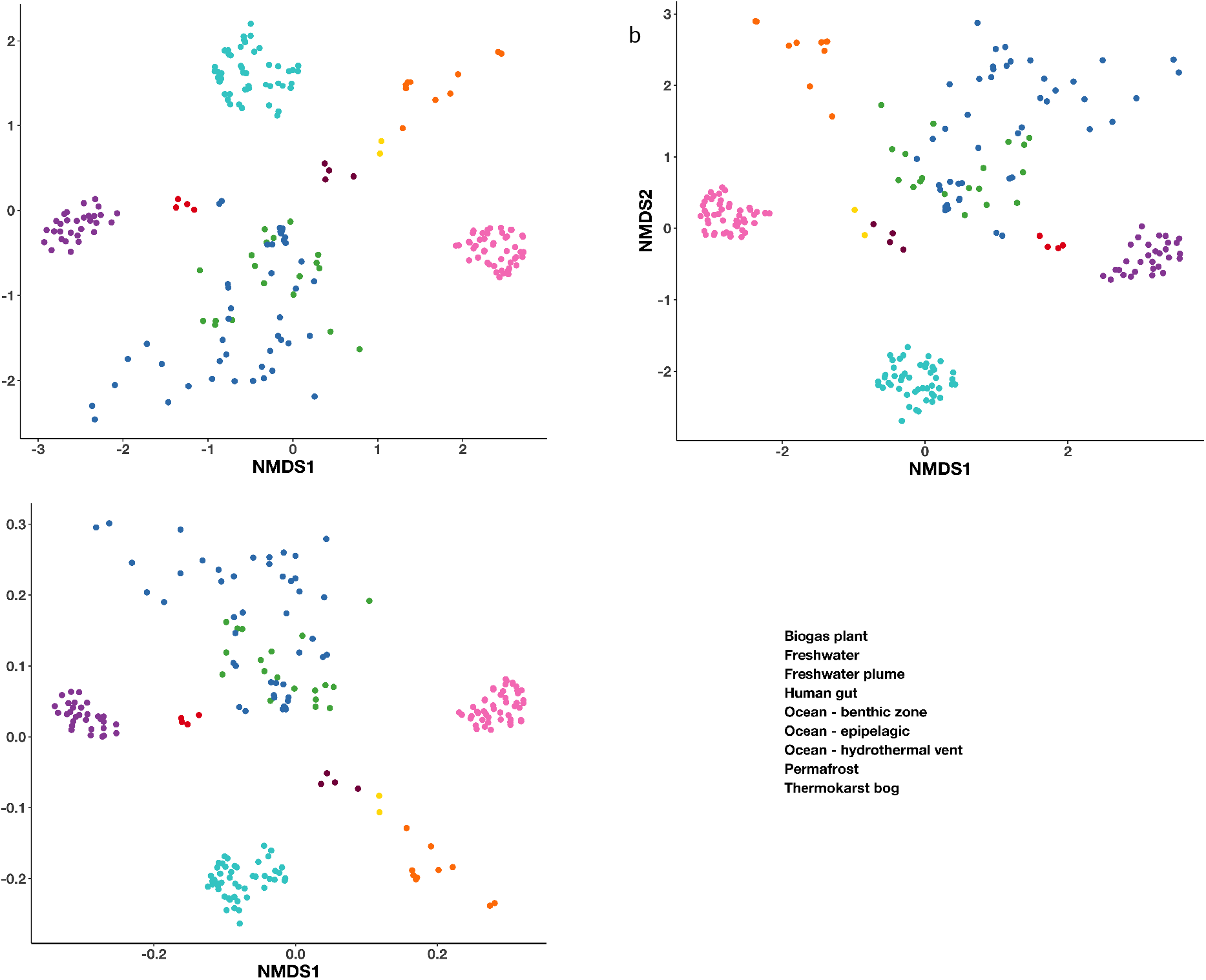
Non-metric multidimensional scaling (NMDS) based ordination of samples by environment. Sample distances are based on Bray-Curtis dissimilarities of a) abundance (% DNA) b) activity (% RNA) c) specific activity (RNA:DNA ratios).

**Figure S8.**
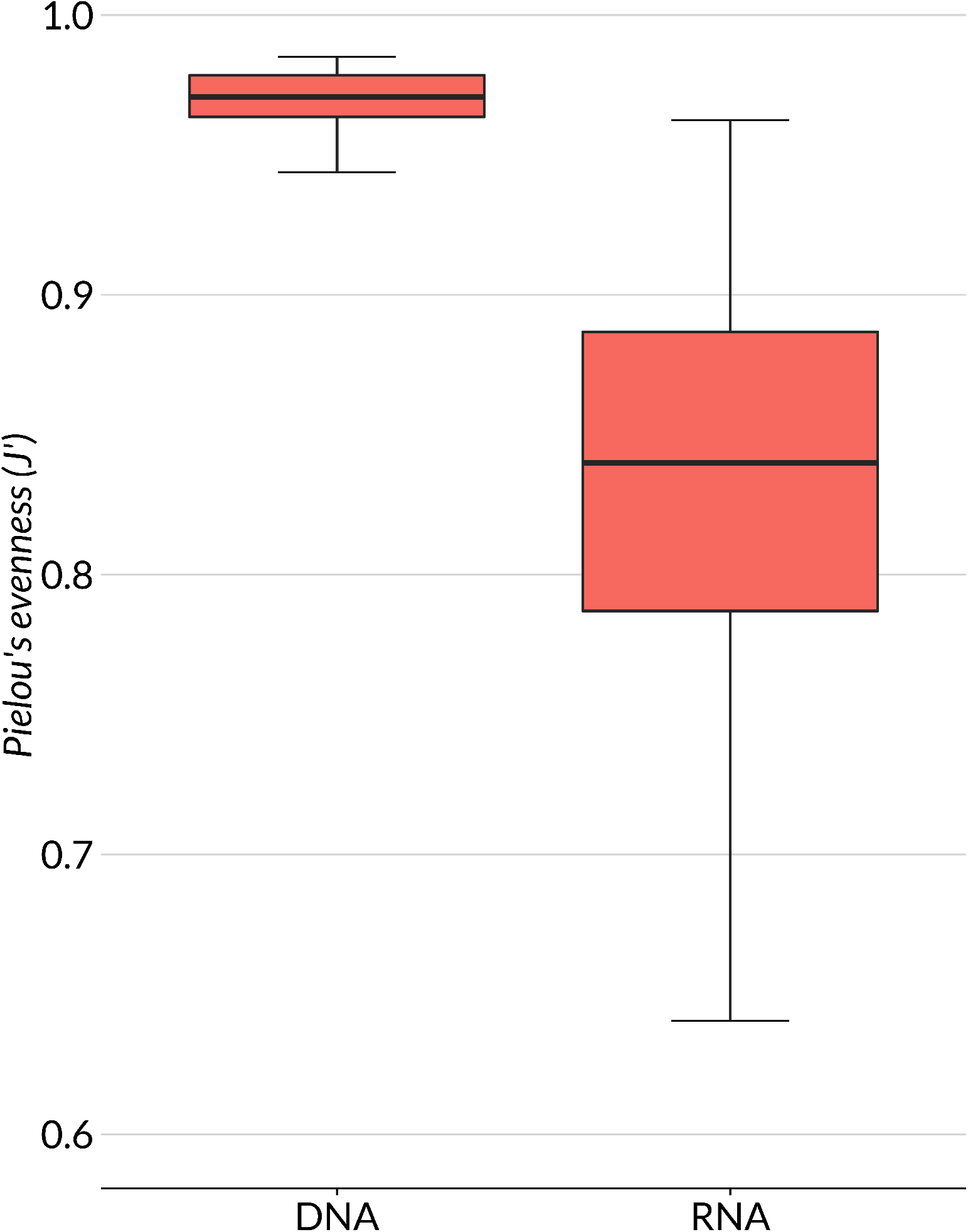
Box and whisker plots of abundance (DNA) and expression (RNA) evenness across all Illumina sampled sequences. Evenness is calculated as Pielou’s evenness (*J*’) of sequence assemblies and their relative frequencies in metagenomes and metatranscriptomes (Mann–Whitney *U* test; *P* < 2.2 x 10^−16^).

**Figure S9.**
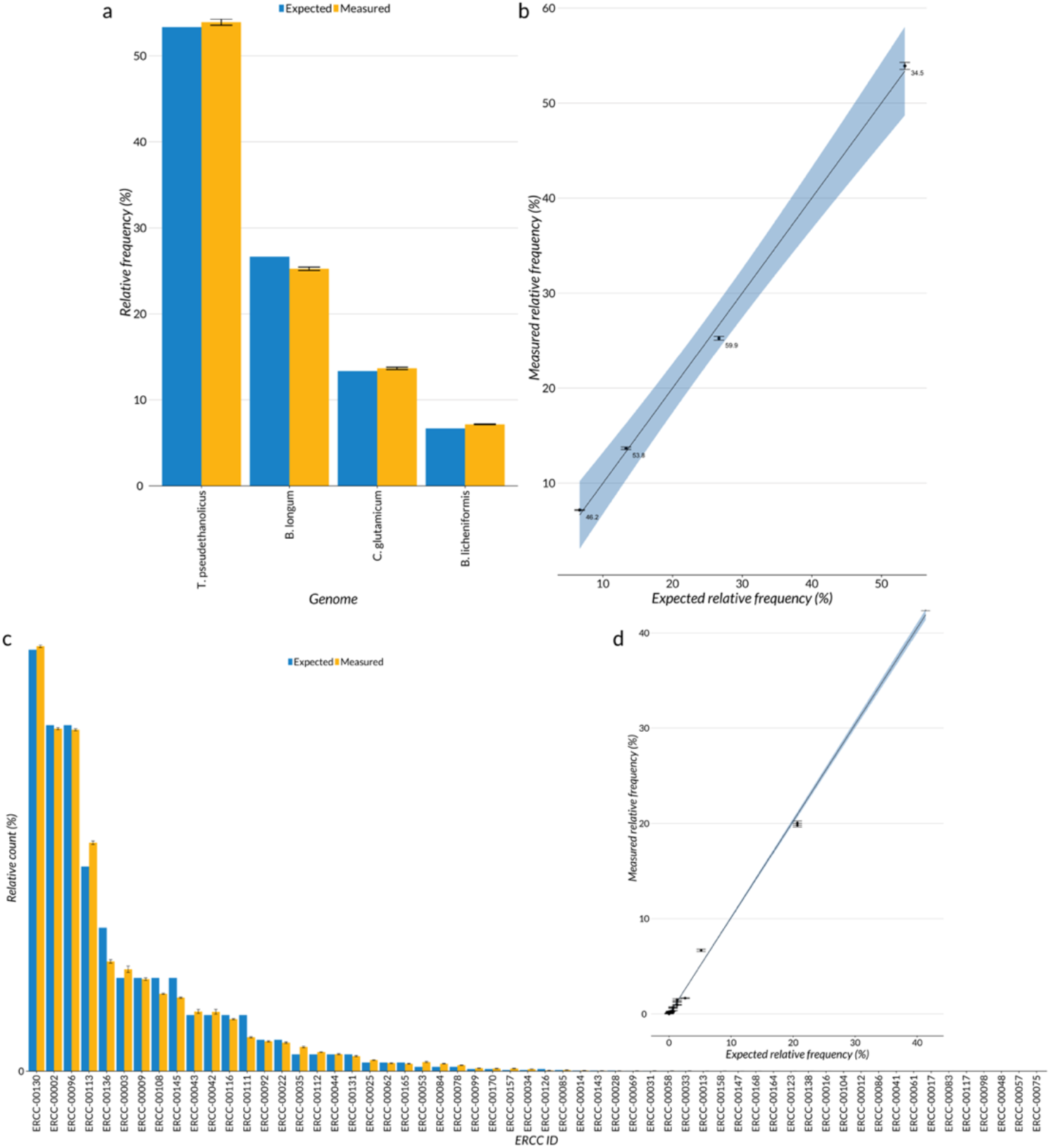
Evaluation of control sequences. a) Mean measured and relative frequencies from SPOT benthic zone metagenomes for each amended genome control with SEM (n = 32). b) Linear regression of measured metagenomic relative frequencies against expected relative frequencies of SPOT benthic zone amended genome controls plotted with 95% confidence interval and SEM (n = 32; R^2^ = 0.9933). c) Mean measured and relative frequencies from SPOT benthic zone metatranscriptomes for each ERCC control with SEM (n = 30). d) Linear regression of measured metatranscriptomic relative frequencies against expected relative frequencies of SPOT benthic zone ERCC controls plotted with 95% confidence interval and SEM (n = 30; R^2^ = 0.9932)

**Table S1. Data sources used in this study (see excel table).** Most data sources are provided by NCBI SRA or ENA accession numbers. Sequence libraries are characterized by nucleic acid source (DNA or RNA), sequencing technology, preparation methods, source environment, attached/free lifestyle, and sample temperature. Pair bin denotes matching DNA and RNA samples. Pairing information was retrieved from the data source databases and source publications. Public data were retrieved from: Alberti et al., 2017^29^, Baker et al., 2012^32^, Bremges et al., 2015^24^, Dupont et al., 2015^26^, Franzosa et al., 2014^21^, Gilbert et al., 2010^27^, Hultman et al., 2015^22^, Li et al., 2015^33^, Maus et al., 2016^23^, Satinsky et al., 2014a^19^, Satinsky et al., 2014b^20^, Satinsky et al., 2015^18^, Shi et al., 2011^25^, Sieradzki et al., 2017^30^, Sunagawa et al., 2015^28^, and Thrash et al., 2017^31^.

**Table S2. RNA contribution of rare and abundant microbes (see excel table).** % RNA of rare and abundant microbes was totaled for each sample sequenced on Illumina platforms. Total % RNA across samples was calculated by summing % RNA for across all samples and renormalizing to the summed % RNA for each abundance fraction.

**Table S3. Annotations of the most expressed ORFs (see excel table).** RefSeq and KEGG annotations of the most expressed ORFs from each sample pairing that had a RefSeq match.

**Table S4.**
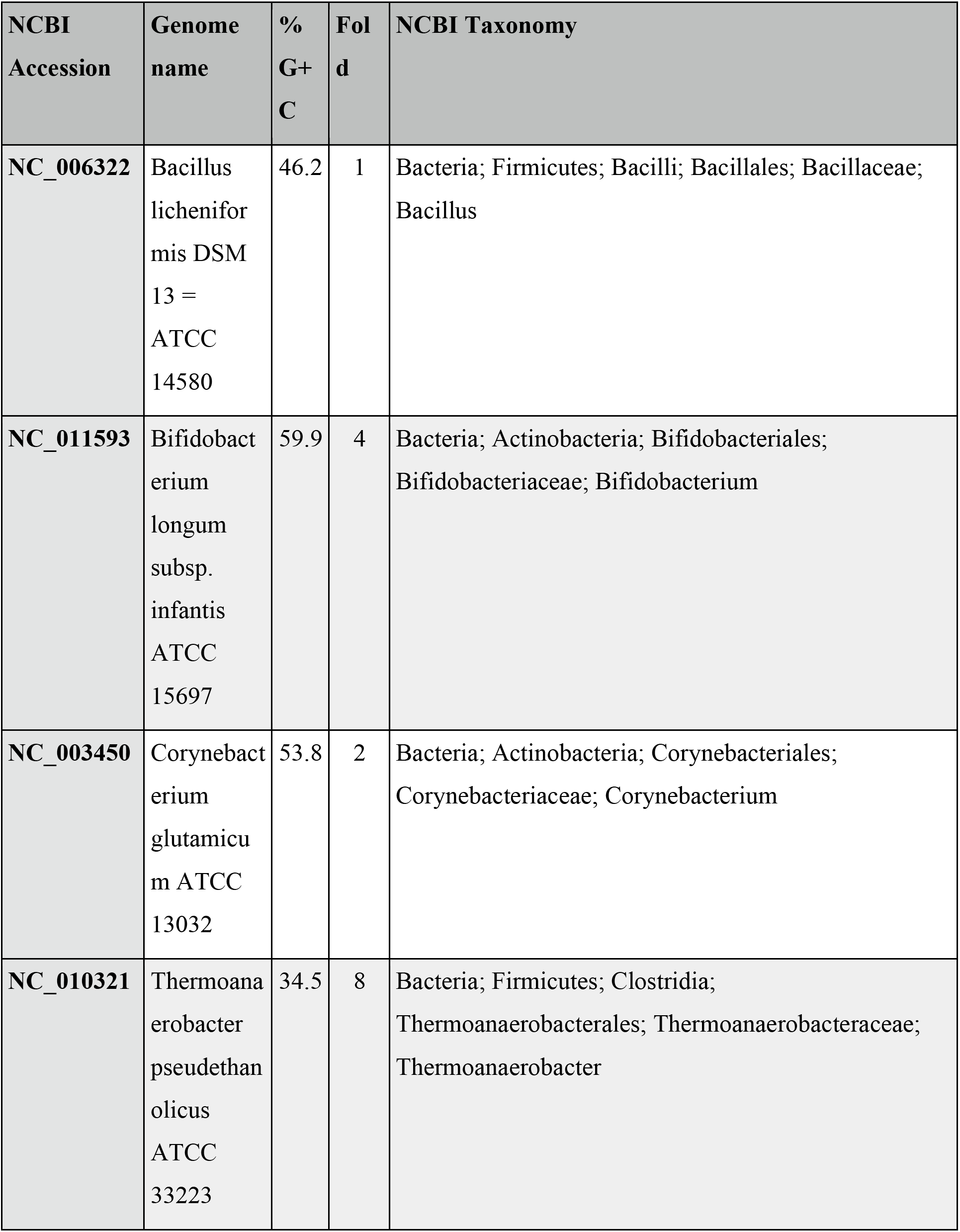
Genome controls used to amend SPOT metagenomes. Each genome was selected to encompass a range of %G+C.

| NCBI Accession | Genome name | % G+C | Fold | NCBI Taxonomy |
| --- | --- | --- | --- | --- |
| NC_006322 | Bacillus licheniformis DSM 13 = ATCC 14580 | 46.2 | 1 | Bacteria; Firmicutes; Bacilli; Bacillales; Bacillaceae; Bacillus |
| NC_011593 | Bifidobacterium longum subsp. infantis ATCC 15697 | 59.9 | 4 | Bacteria; Actinobacteria; Bifidobacteriales; Bifidobacteriaceae; Bifidobacterium |
| NC_003450 | Corynebacterium glutamicum ATCC 13032 | 53.8 | 2 | Bacteria; Actinobacteria; Corynebacteriales; Corynebacteriaceae; Corynebacterium |
| NC_010321 | Thermoanaerobacter pseudethanolicus ATCC 33223 | 34.5 | 8 | Bacteria; Firmicutes; Clostridia; Thermoanaerobacterales; Thermoanaerobacteraceae; Thermoanaerobacter |

